# miR-495-3p Sensitizes BCR::ABL1 Expressing Leukemic cells to Tyrosine Kinase Inhibitors by Targeting Multidrug Resistance 1 Gene including in T315I Mutated cells

**DOI:** 10.1101/2022.10.17.512501

**Authors:** Yutthana Rittavee, Jérôme Artus, Christophe Desterke, Isidora Simanic, Lucas Eduardo Botelho de Souza, Sandra Riccaldi, Sabrina Coignard, Yousef Ijjeh, Patricia Hughes, Annelise Bennaceur-Griscelli, Ali G. Turhan, Adlen Foudi

## Abstract

Chronic myeloid leukemia (CML) is a clonal hematopoietic malignancy driven by the BCR::ABL1 fusion oncoprotein. The development of tyrosine kinase inhibitors (TKIs) has deeply increased long-term survival of CML patients. Nonetheless, one patient out of four will switch TKI off owing either to drug intolerance or resistance partly due to amplification or mutations of *BCR::ABL1* oncogene and alteration of ATP-binding cassette (ABC) transporters. Increasing evidence suggests an involvement of the microRNA miR-495-3p in cancer-associated chemo-resistance through *multidrug resistance 1* (*MDR1*) gene which encodes an ATP-dependent efflux pump. Our study aimed at investigating the potential role of miR-495-3p in CML TKI chemo-sensitivity and determining the underlying molecular circuitry involved. We first observed that *miR-495-3p* expression was lower in BCR::ABL1 expressing cellular models *in vitro*. Notably, loss-of-function experiments showed increased proliferation associated with a decreased number of non-dividing cells (G0/G1) and resistance to Imatinib. Conversely, our data showed that *miR-495-3p* overexpression hindered leukemic cell growth and TKI resistance even in Imatinib-resistant T315I-mutant cells as well as drug efflux activity through *MDR1* regulation. To further investigate the role of miR-495-3p in CML patients, we found that predicted miR-495-3p targets were upregulated in patients in blast crisis involved in protein phosphorylation and associated with the worst prognosis. Taken together, our results demonstrate that down-regulation of *miR-495-3p* expression is important in the malignant phenotype of CML and TKI resistance mechanisms, which could be a useful biomarker and a potential therapeutic target to eradicate CML.

**MeSH terms:**

- ATP Binding Cassette Transporter, Subfamily B
- ATP Binding Cassette Transporter, Subfamily B, Member 1 / drug effects
- ATP Binding Cassette Transporter, Subfamily B, Member 1 / metabolism*
- Blast Crisis / pathology
- Cell Line, Tumor
- Cell Proliferation / drug effects*
- Cell Survival / drug effects
- Drug Resistance
- Genes, MDR
- Imatinib Mesylate
- Leukemia, Myelogenous, Chronic, BCR-ABL Positive
- MicroRNAs / genetics
- MicroRNAs / physiology*

**HIGHLIGHTS:**

- miR-495-3p inhibits leukemic cell growth and is downregulated in BCR::ABL1 expressing cell lines
- miR-495-3p modulates response to TKI treatment including in UT7 cells expressing T315I
- Overexpression of miR-495-3p leads to a decrease of *MDR1* and drug efflux activity
- Bioinformatics analyses reveal that MiR-495-3p target genes are upregulated in blast crisis

## INTRODUCTION

Chronic myeloid leukemia (CML) is a clonal malignant myeloproliferative neoplasm characterized by the uncontrolled proliferation of myeloid cells. Primarily, BCR::ABL1 acts as a constitutively active tyrosine kinase that alters cell proliferation, differentiation, survival, and genetic stability in hematopoietic cells in which it is expressed [1,2].

CML therapy has greatly benefited from the fact that BCR::ABL1 expressing cells exhibit the oncogene-dependence phenomenon such that BCR::ABL1 inhibition has been found to lead to the inhibition of leukemic cell growth [3]. Imatinib (IM) treatment has been the first TKI licensed for clinical use after its validation in preclinical experimental models and the first clinical trial IRIS comparing IM to the standard therapy IFN-ARA-C [4]. It has improved deep molecular responses in more than 20-30% of newly diagnosed chronic CML patients [5]. However, several works have now shown that CML stem cells are resistant to TKI, due to their quiescence and their oncogene independence [6–8]. Moreover, responses to TKI are altered by either intolerance or by the occurrence of several resistance phenomena which have been explored in detail, including decreased OCT1 expression responsible for the entry of IM into the cells and increased expression of MDR1 [9]. The resistance is also due in 10-20% of cases to mutations in the ABL-kinase domain which impair the accessibility of TKI drugs to the ATP pocket either by direct interaction or by conformational changes. The development of resistance to TKI treatment which occurs in CML patients severely impairs the clinical application but led to the development of novel TKI of the second generation [10–12] and more recently, the development of the Asciminib, able to inhibit BCR::ABL1 TKI activity by binding to the myristoyl binding pocket [13].

The ‘gatekeeper mutation’ T315I mutation in the Imatinib-binding site is the most common mutation amongst BCR::ABL1-dependent resistance, identified in 4–15 percent of patients with IM resistance, and is a major concern because it renders leukemic cells resistant to all licensed TKI except Ponatinib [14]. Meanwhile, TKI resistance can also occur independently of BCR::ABL1. The activity of drug transporters is one of the mechanisms that has been identified as mediating TKI resistance [15,16]. The P-glycoprotein (P-gp or MDR1 protein), an ATP-binding Cassette (ABC) efflux transporter encoded by the human multidrug-resistant *ABCB1/MDR1* gene [17], encodes key IM efflux pumps and has been proposed as the main drug resistance mediator in CML [9,18]. Failure modes of TKI monotherapies have prompted the exploration of the mechanisms regulating drug resistance. On that account, multiple microRNAs have been implicated in the drug resistance of CML [19,20].

MicroRNAs are noncoding, single-stranded RNAs with 18 to 25 nucleotides that act as epigenetic negative regulators of specific gene targets at the post-transcriptional and translational levels by binding to the 3′-untranslated region (3′-UTR) of targeted mRNAs; causing in translational repression and/or mRNA degradation. Over recent years, the dysregulation of microRNA expression has been increasingly considered a regulator involved in the progression and the chemoresistance of various cancer types, including CML [21,22]. *microRNA-495-3p* (*miR-495-3p*) is a member of the 14q32.31 microRNA cluster. Accumulating evidence has linked *miR-495-3p* deregulation to cancer progression [23]. miR-495-3p was shown to directly regulate MDR1 translation, which reduces drug efflux from cancer cells [24]. Notably, Xu et al. discovered that drug-resistant CML K562 cell line expressed lower levels of *miR-495-3p* and that transfecting those cells with *miR-495-3p* mimics restores their susceptibility to Adriamycin chemotherapeutic agent [25]. Still, the exact role of miR-495-3p in drug resistance to TKIs in CML remains unknown.

In this study, we evaluated the expression of *miR-495-3p* expression in leukemic cell models expressing BCR::ABL1 either constitutively or in a DOX-inducible context. We have found that there is a BCR::ABL1-induced down-regulation of miR-495-3p allowing us to investigate the role of this phenomenon in TKI resistance including in cells expressing T315I.

## MATERIAL AND METHODS

### Cell culture and generation of stable cell lines

The UT-7 which is a GM-CSF dependent human hematopoietic cell line [26] and the KCL22 Ph1 chromosome-positive CML cell line [27] were cultured in Roswell Park Memorial Institute-1640 (RPMI-1640) medium (Gibco, Rockville, MD, USA) [containing 10% fetal bovine serum (FBS), 1% penicillin-streptomycin, and 10 ng/mL granulocyte/macrophage colony-stimulating factor (GM-CSF) for UT-7] under an atmospheric condition of 5% CO_2_ at 37 °C. BCR::ABL1 positive variants of UT-7 cell line were obtained from previous studies including UT7-11 expressing wild-type BCR::ABL1 [28], UT-7 BCR::ABL1^ind^ expressing TET-OFF-inducible BCR::ABL1 [29] and UT7-T315I expressing T315I-mutated version of BCR::ABL1 [30]; which were cultured under the same conditions as mentioned above. The medium was subsequently replaced with fresh medium every 2 days.

For gain- and loss-of function studies, we used the following vectors: miR-495-3p (HmiR0226-MR03), AntagomiR-495-3p (MmiR-AN0537-AM04), and control vectors (CmiR0001-MR03 and CmiR-AN0001-AM04, respectively) all from GeneCopoeia (Rockville, MD, USA). Lentiviral particles were produced and used for transduction at a MOI of 1. Stable cells expressing fluorescent reporter proteins (mCherry or eGFP) were sorted using BD FACSAria Flow Cytometer (BD Bioscience, Mountain View, CA, USA).

### RNA extraction and quantitative real-time PCR

Total RNAs were extracted using the PureLink® RNA Mini Kit (Invitrogen, Carlsbad, CA, USA) and used for reverse transcription using SuperScript™ III Reverse Transcriptase (Invitrogen) with random primers or a universal stem-loop primer for microRNA detection as described in [31]. Real-time PCR was carried out on Agilent Mx3005P QPCR System (Agilent Technologies, Santa Clara, CA, USA) with the FastStart Universal SYBR Green Master (Roche, Mannheim, Germany). Primers sequences were: *miR-495-3p*, F: 5’-GCCAAACAAACATGGTGCACTTC-3’, R: 5’-CGAGGAAGAAGACGGAAGAAT-3’; *U6*, F: 5’-GCTTCGGCAGCACATATACTAAAT-3’, R: 5’-CGAGGAAGAAGACGGAAGAAT-3’; *MDR1*, F: 5’-CCCATCATTGCAATAGCAGG-3’, R: 5’-GTTCAAACTTCTGCTCCTGA-3’; *B2M*, F:5’-TGCTGTCTCCATGTTTGATGTATCT-3’, R: 5’-TCTCTGCTCCCCACCTCTAAGT-3’.

### Cell proliferation, colony formation (CFC), cell cycle, and cell viability assays

For cell proliferation assays, 5×10^3^ UT7 cells were plated per well of a 96-well plate and counted every two days (in three replicates) using trypan blue exclusion.

For clonogenic assays of UT7 cells and their BCR::ABL expressing counterparts, 500 cells were seeded in 35-mm dish in methylcellulose media (H4435, Stem Cell Technologies, Vancouver, Canada). Colonies were photographed and scored 14 days later using ImageJ software (LOCI, University of Wisconsin, Madison, USA).

Cell cycle analysis was performed using Hoechst 33342 and BD LSR Fortessa Flow Cytometer (BD Bioscience). Data were analyzed using FlowJo v10.1 software (FlowJo LLC, Ashland, OR, USA).

Cell viability was determined using CyQUANT™ XTT Cell Viability Assay (Thermo Fisher Scientific, Waltham, MA, USA) according to the manufacturer’s protocol. 5×10^3^ cells were plated per well of a 96-well and cultured with or without 1 μM Imatinib or 100 nM Ponatinib for 48 h. After 4h staining with XTT mixture, absorbances were measured using a Modulus™ II Microplate Reader (Turner BioSystems, Sunnyvale, CA, USA) at 450 and 600 nm.

### MDR1-mediated drug efflux activity assay

Drug efflux activity was determined by intracellular accumulation of Rhodamine 6G (Rho6G) (Sigma-Aldrich, France). Cells were cultured with 10 μM Rho6G for 30 min at 37 °C in the dark, washed 3 times with ice-cold media, transferred to fresh pre-warmed Rho6G-free media, and subsequently cultured at 37 °C. A similar amount of cells was collected in ice-cold media every 5 min for time-course experiments. Cells were maintained on ice and fluorescence imaging and intensity measurements were performed using a Modulus™ II Microplate Reader at 450 nm. The interference of GFP expression was normalized by unstained samples.

### Statistical analysis

All the experiments were repeated three times independently. All data were analyzed with GraphPad Prism 8.4.3 (GraphPad Software, San Diego, CA, USA) and represented as the mean ± standard deviation (SD). The differences between two groups were evaluated using unpaired two-tailed Student’s t-tests. Differences for multiple comparisons were subjected to one-way analysis of variance followed by Dunnett’s test. A p-value of less than 0.05 was considered to be statistically significant.

### CML transcriptome dataset

Normalized matrix of Transcriptome dataset GSE4170 [32] was downloaded at the following address: https://www.ncbi.nlm.nih.gov/geo/query/acc.cgi?acc=GSE4170 (accessed on 2022, April 22nd) and annotated with GPL2029 Rosetta/Merck Human 25k v2.2.1 microarray technology platform available at the address: https://www.ncbi.nlm.nih.gov/geo/query/acc.cgi?acc=GPL2029 (accessed on 2022, April 22nd). The resulting gene annotated matrix was used for downstream bioinformatics analyses.

### RNA-sequencing TCGA AML cohort

Through the Cbioportal web application tool [33], clinical and RNA-sequencing Z-score V2 data were aggregated for all diploid samples inside the TCGA AML cohort [34]. These data concerning experiments were performed on 173 *de novo* AML samples. Among this AML cohort, 46 percent of patients present a normal karyotype, and 12 percent of them present a complex karyotype. Age median diagnosis was 57 years old for this AML cohort.

### Transcriptome analyses

Prediction of mRNA targets of hsa-miRNA-495-3p was done in two distinct databases: miRDB [35] and targetscan [36]. Inner join between was done to build a Venn diagram and common target lists obtained from the two databases were retained to perform transcriptome analyses in the GSE4710 dataset. Bioinformatics analyses were performed in R software environment version 4.1.3. Unsupervised principal component analysis was done with “prcomp” base function and output plot was drawn with “autoplot function” of ggfortify R-package version 0.4.14. Differential expressed gene analysis was done with a linear model for microarray algorithm implemented in limma R-package version 3.48.3 [37]. Expression heatmap with unsupervised hierarchical clustering was down with pheatmap R-package version 1.0.12 based on Euclidean distances and Ward.D2 method. Protein-protein interaction network was built with online proteomic tool NetworkAnalyst [38] with an inference of Gene Ontology Biological Process (GO-BP) database [39].

## RESULTS

### *miR-495-3p* is downregulated in BCR::ABL-expressing cell lines and regulates cell proliferation and IM resistance

To investigate a possible role of miR-495-3p in CML, we first analyzed *miR-495-3p* expression in various cell lines including KCL22, BCR::ABL1-expressing UT-7/11 and the parental UT-7. Interestingly, compared to UT-7, *miR-495-3p* expression was decreased in KCL22 and in UT-7/11 (Figure 1A). To confirm this phenomenon, we used our TET-OFF inducible system UT/7 cell line where *BCR::ABL1* is expressed in the absence of doxycycline. In these cells, *miR-495-3p* expression was also decreased in the absence of doxycycline and so when BCR::ABL1 was expressed (Figure 1B). Taken together, *miR-495-3p* expression appears to be inversely correlated to *BCR::ABL1* expression at the transcriptional level, suggesting a possible role of miR-495-3p in BCR::ABL-induced leukemogenesis.

**Figure 1:**
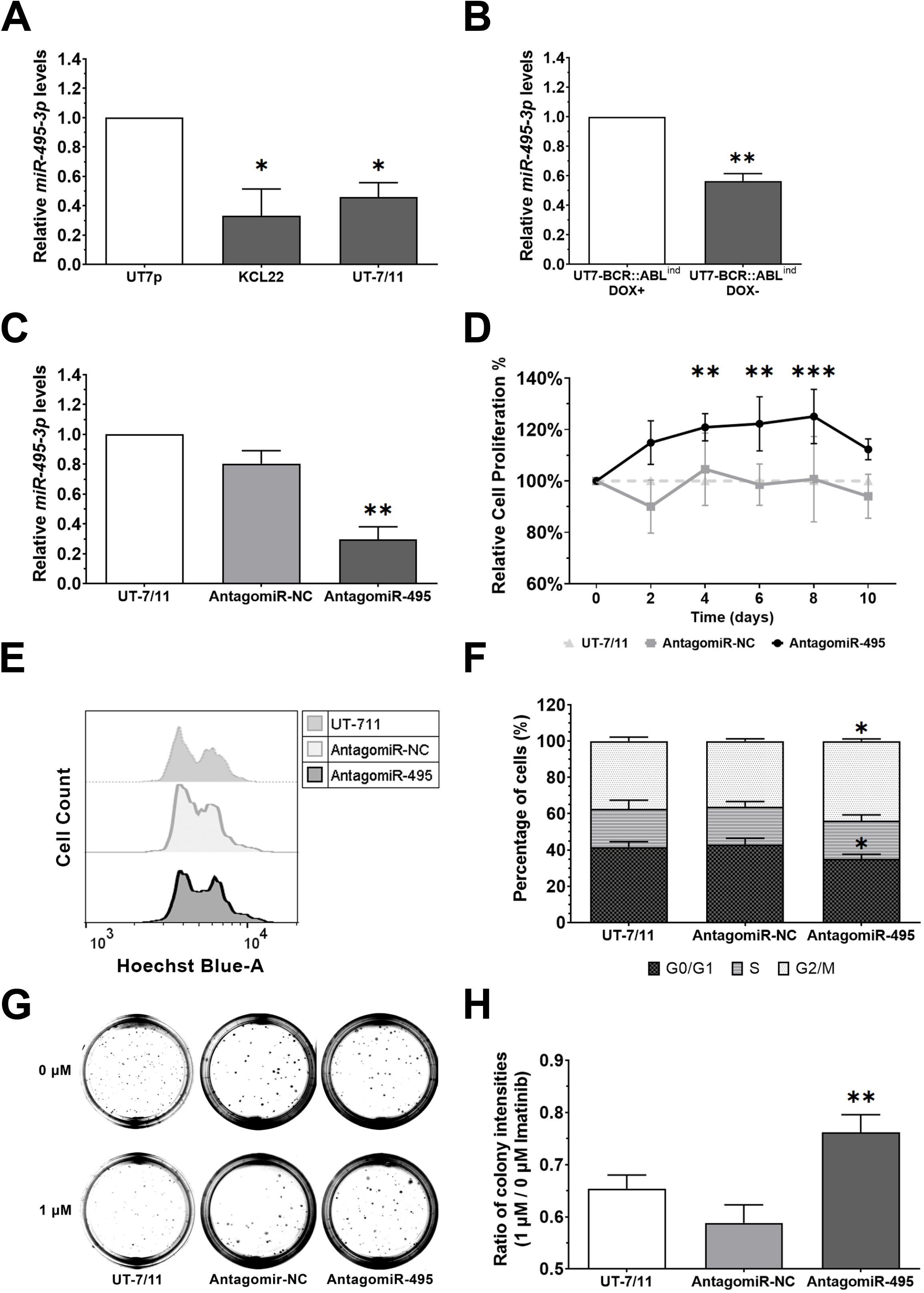
Expression of *miR-495-3p* and effects of *miR-495-3p* knock-down in CML cell lines. **(A-C)** Relative quantification of *miR-495-3p* expression by real-time PCR normalized to *U6* expression. (A) *miR-495-3p* expression in KCL22 (fold change (FC) =−3.01 ± 0.18, p = 0.0185) and UT-7/11 (FC = −2.17 ± 0.10, p = 0.0329) normalized to UT7p. (B) *miR-495-3p expression* in BCR::ABL1-inducible UT7 cell lines in the presence (DOX+) or absence (DOX-) of doxycycline (FC = −1.77 ± 0.05, p = 0.044). (C) *miR-495-3p* expression in UT-7/11 cells stably expressing AntagomiR-495-3p or its control vector (antagomiR-NC) (FC = −3.38 ± 0.09, p = 0.0076). **(D)** Cell proliferation of UT-7/11 cells expressing the empty vector AntagomiR-NC or AntagomiR-495-3p relative to UT-7/11 (p = 0.0054 on day 4, p = 0.0030 on day 6, and p = 0.0008 on day 8). Cells were plated at equal density and cultured for 10 days. Cell numbers were determined using trypan blue exclusion. **(E)** Cell cycle analysis using flow cytometry quantification of DNA content with Hoechst 33342 dye (G0/G1 phase; 34.98% *versus* 41.63% in control; p = 0.0345, G2/M phase; 43.87% *versus* 37.47% in control; p = 0.0425). **(F)** Quantitative analysis of cell cycle phase distribution of UT-7/11 expressing the AntagomiR-NC or AntagomiR-495-3p. **(G)** Effect of *miR-495-3p* overexpression in CFC assays. Representative pictures of cell culture dishes after 2 weeks in culture of UT-7/11 cells expressing AntagomiR-NC or AntagomiR-495 in the absence (0 μM) or presence (1 µM) of IM. **(H)** Quantitative analysis of CFC assays. The bar chart shows the ratio between colony numbers with IM and without. Each data represents mean ± SD from three independent experiments (mean ratio = 0.76 ± 0.03 *versus* 0.65 ± 0.03 in UT7/11 cells, p=0.0096). The One-way or Two-way ANOVA approach was used to assess the statistical significance. *, p< 0.05; **, p < 0.01; ***, p < 0.001.

To test this hypothesis, we developed a loss-of-function strategy using an AntagomiR-495-3p. We first validated its functionality by confirming that AntagomiR-495 reduces *miR-495-3p* expression contrary to the control Antagomir-NC (Figure 1C). Strikingly, we observed that miR-495-3p inhibition markedly promoted UT7/11 cell proliferation compared to Antagomir-NC (Figure 1D) which was associated with a decrease in the proportion of cells in G0/G1 phase but an increase in G2/M phase (Figure 1E, 1F). We next tested whether miR-495-3p inhibition could also affect the sensitivity of CML cells to TKI treatment. To do this, we measured the ability of UT7/11 cells expressing AntagomiR-NC or AntagomiR-495 to form colonies in the presence of Imatinib. We first observed that IM treatment impaired colony formation in all cell lines (Figure 1G, 1H). Surprisingly, its toxic effect was strongly reduced in the presence of AntagomiR-495. Collectively, these results demonstrate that *miR-495-3p* knockdown promotes cell proliferation and IM resistance in BCR::ABL1 expressing cells.

### Overexpression of *miR-495-3p* decreases leukemia cell proliferation but increases IM sensitivity

To further characterize its role, we undertook gain-of-function experiments in which *miR-495-3p* was overexpressed in UT-7/11 following lentiviral transduction (Figure 2A). We noticed that miR-495-UT-7/11 cells exhibited a reduced proliferation capacity associated with an increase in the proportion of cells in G0/G1 and a reduction in G2/M phase (Figure 2B, 2C). Importantly, these phenotypes were not observed in NC-UT-7/11 cells expressing a control scrambled sequence. We next tested the impact of *miR-495-3p* overexpression on IM sensitivity. As expected, IM strongly decreased UT-7/11 and NC-UT-7/11 cell viability (Figure 2D). Cell viability was even more reduced in miR-495-UT-7/11 cells indicating that miR-495-3p overexpression enhances cell sensitivity to TKI treatment. To further validate this observation, we analyzed the effect of IM treatment in clonogenic assays using UT/7 cells. In all cases, IM treatment reduced colony formation but this reduction was exacerbated in miR-495-UT-7/11 compared to UT-7/11 and NC-UT-7/11 cells (Figure 2E, 2F). Taken together, complementary gain- and loss-of-function experiments demonstrate that *miR-495-3p* is a critical negative regulator of cell cycle progression and IM resistance.

**Figure 2:**
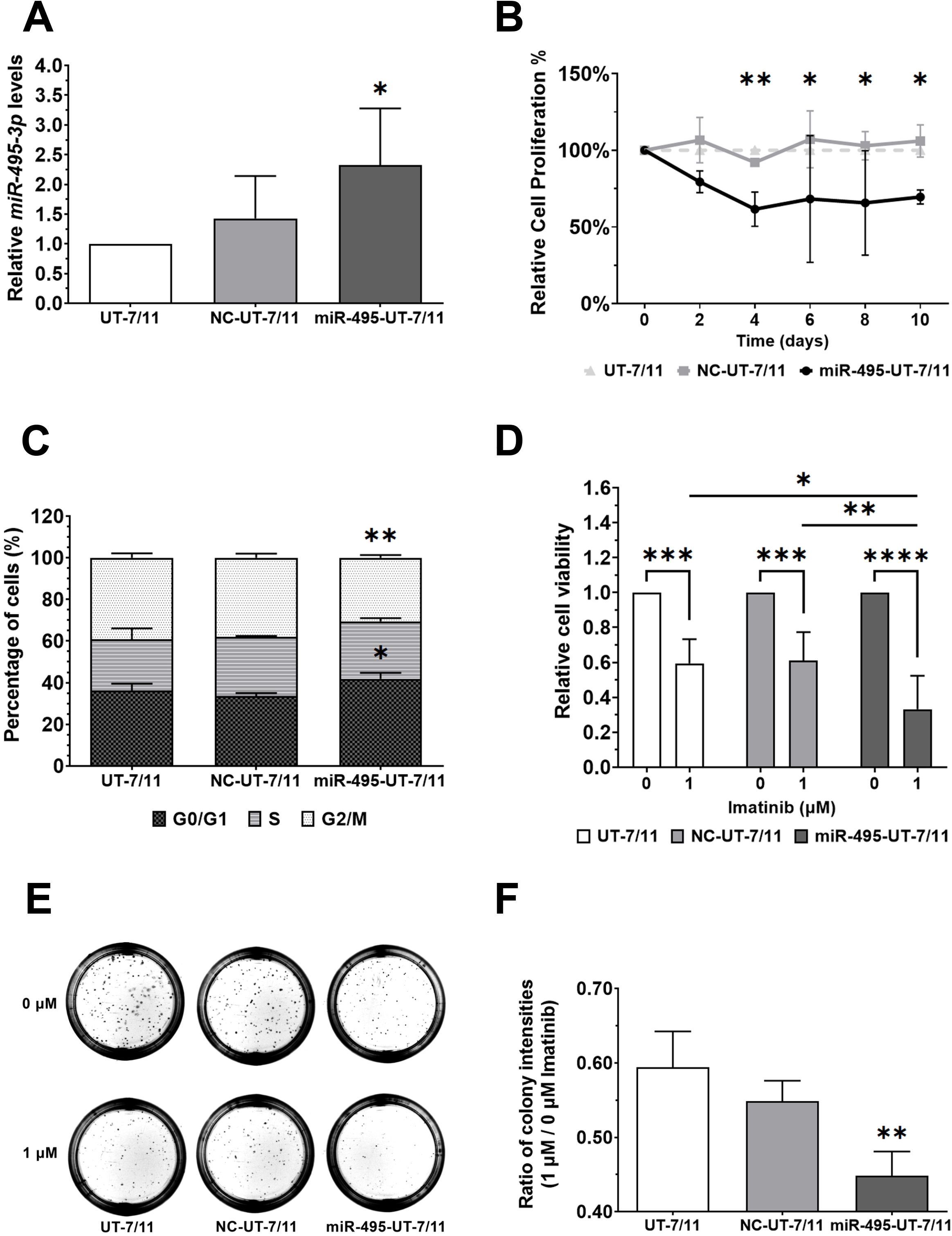
Effect of *miR-495-3p* overexpression on UT-7/11 cell proliferation and IM resistance. **(A)** Relative quantification of *miR-495-3p* expression by real-time PCR in UT-7/11 cells stably expressing miR-495-3p (miR-495-UT-7/11) or its control vector (NC-UT-7/11) (FC = 2.32 ± 0.95, p = 0.0472). *miR-495-3p* expression was normalized to *U6* expression. **(B)** Cell proliferation of miR-UT7/11 and NC-UT-7/11 relative to UT-7/11. Cells were plated at equal density and cultured for 10 days (p = 0.0072 on day 4, p = 0.0298 on day 6, p = 0.0176 on day 8, and p = 0.0380 on day 10). Cell numbers were established using trypan blue exclusion assay. **(C)** Quantitative analysis of the cell cycle phase distribution (G0/G1 phase; 41.94% *versus* 36.24% in UT-7/11; p = 0.0286, G2/M phase; 30.47% *versus* 39.19% in UT-7/11; p = 0.0013). **(D)** Cytotoxic effects of IM were measured using XTT assay (40.58% reduction in UT-7/11, 39.04% in NC-UT-7/11, and 66.77% in miR-495-UT-7/11, p=0.0008 compared to UT-7/11 and p=0.0005 compared to NC-UT-7/11). **(E)** Effect of *miR-495-3p* overexpression in CFC assays. Representative pictures of cell culture dishes after 2 weeks in culture of UT-7/11, NC-UT7/11, and miR-495-UT-7/11 cells in the absence (0 μM) or presence (1 µM) of IM. **(F)** Quantitative analysis of CFC assays. The bar chart shows the ratio between colony numbers obtained in the presence and absence of IM (mean ratio = 0.45 ± 0.03 *versus* 0.59 ± 0.05 in UT7/11 cells, p=0.0072). Each data represents mean ± SD from three independent experiments. The One-way or Two-way ANOVA approach was used to assess the statistical significance. *, p < 0.05; **, p < 0.01; ***, p < 0.001; ****, p < 0.0001.

### *miR-495-3p* negatively regulates the expression and efflux activity of MDR1

*miR-495-3p* plays multiple roles in physiological and pathological situations including cell proliferation and drug resistance in multiple cancers [24]. Particularly, according to the DIANA-microT database [40], *miR-495-3p* is predicted to bind to 2 binding sites localized in the 3′-UTR of *MDR1* with a high microRNA targeted genes (miTG) score of 0.7528 (Figure 3A). To test whether the regulation of IM resistance by *miR-495-3p* is mediated by MDR1, we first quantified *MDR1* expression and observed a significantly lower expression in miR-495-3p overexpressing cells compared to UT-7/11 cells (Figure 3B). These data suggested that miR-495-3p might regulate *MDR1* expression in BCR::ABL1 expressing cells.

**Figure 3:**
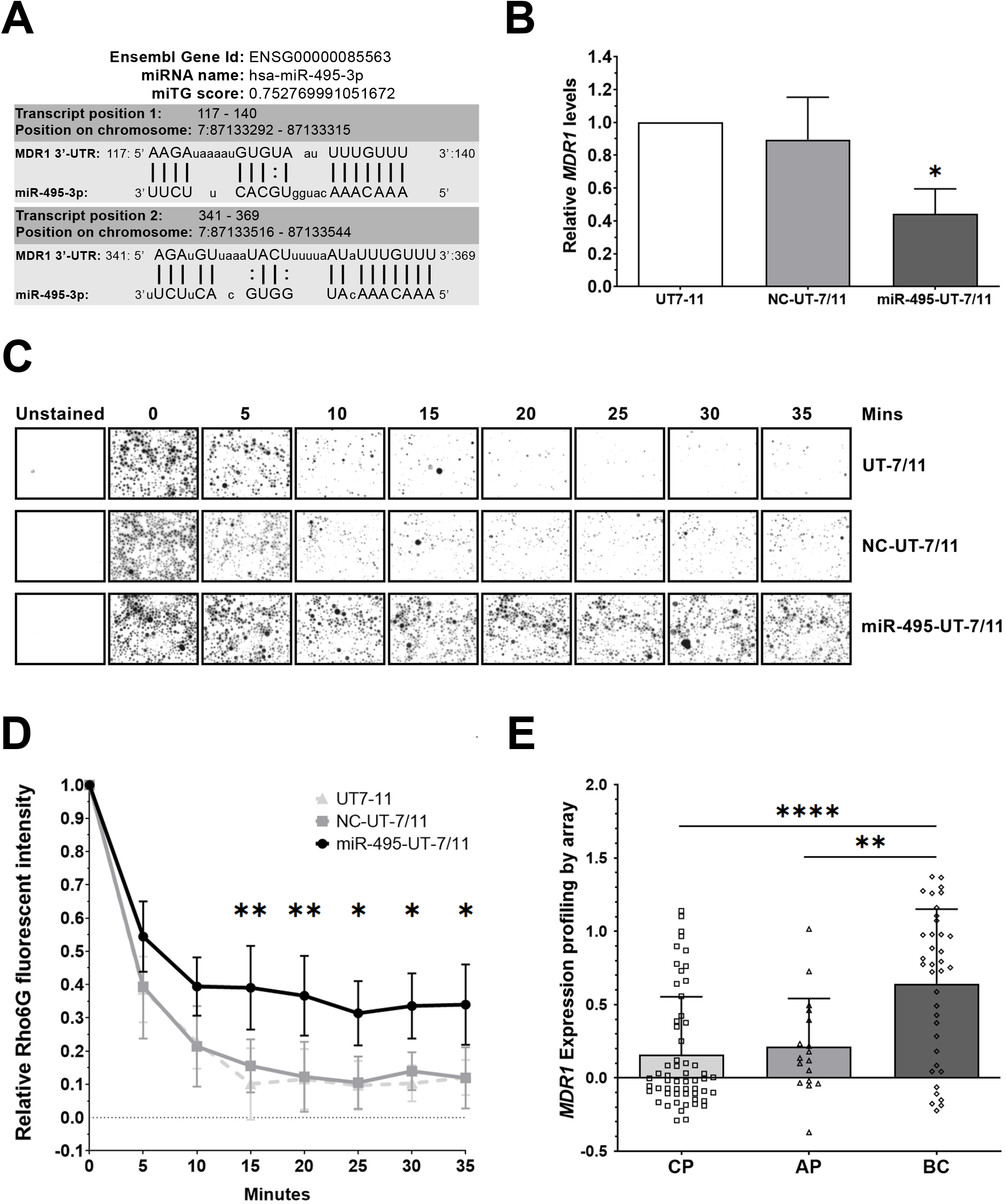
Effect of miR-495-3p on *MDR1* gene expression and efflux activity. **(A)** Alignment between the miRANDA-predicted miR-495-3p sequence and the *MDR1* 3′-UTR WT target sequence. **(B)** Relative quantification of *MDR1* expression by real-time PCR in UT-7/11, NC-UT-7/11, and miR-495-UT-7/11 cells (FC = −2.27 ± 0.09, p = 0.0440). *MDR1* expression was normalized to *B2M* expression. **(C)** Representative images of Rhodamine 6G accumulation in UT-7/11, NC-UT-7/11, and miR-495-UT-7/11 cells. Cells were treated with 1h pulse of 10 µM Rho6G and chased in Rho6G-free media for the indicated periods of time. **(D)** Relative Rho6G fluorescent intensity. Results were normalized to the time point of 0 minutes after washing. Each data represents mean ± SD from three independent experiments. **(E)** *MDR1* mRNA expression profiling by array in patients at different stages of disease, on the basis of RNA-seq data from the GSE4170 transcriptome dataset (0.6429 in BC versus 0.1591 in CP, p=<0.0001; versus 0.2122 in AP, p=0.0024). CP: n = 57, AP: n = 17, BC: n = 37 CML patients. The One-way or Two-way ANOVA approach was used to assess the significance. *, p < 0.05; **, p < 0.01; ****, p < 0.0001.

Next, we analyzed MDR1 activity using a Rhodamine 6G (Rho6G) efflux assay. After a pulse of Rho6G, which is a substrate of MDR1 transporter, we monitored the dye fluorescence dynamics as a measure of the efflux activity in miR-495-3p-overexpressed UT-7/11 cells. We observed that the fluorescence activity was reduced by almost 90% after a chase of 15 to 20 minutes in UT-7/11 and NC-UT-7/11 but remains significantly higher after 15 minutes in *miR-495-3p* overexpressing cells (Figure 3C, 3D). These results suggested that *MDR1* expression might be linked to BCR::ABL1-associated leukemogenesis and TKI resistance as this has previously been reported in primary CML [9,16]. Thus, we analyzed *MDR1* expression in bone marrow samples of patients in chronic (CP), accelerated (AP), and blast crisis (BC) phases (Figure 3E). Interestingly, *MDR1* expression was upregulated in BC compared to CP and AP. Altogether, we propose that miR-495-3p is a negative regulator of IM resistance by suppressing *MDR1* expression.

### Overexpression of *miR-495-3p* suppresses the drug resistance phenotype in Imatinib-resistant CML cell line

Having established the role of miR-495-3p in the regulation of IM resistance, we next tested whether this role could be extended to other TKIs in a drug resistance context. We used the UT7-T315I cell line which expresses the T315I-mutated version of BCR::ABL1 and is resistant to most of the TKIs except Ponatinib [30]. We generated UT7-T315I overexpressing miR-495-3p (Figure 4A) and measured cell viability following IM and Ponatinib treatments. As expected, we observed that IM did not alter cell viability and *miR-495-3p* overexpression had no effect on IM resistance (Figure 4B). However, Ponatinib treatment significantly reduced cell viability of UT7-T315I and NC-T315I UT7 and its effect was even more pronounced in miR-495-T315I cells (Figure 4C). Similarly, we analyzed the effect of Ponatinib in CFC assays. These experiments showed a statistically significant reduction of clonogenic activity in miR-495-expressing UT7-T315I cells compared to UT7 T315I and NC-T315I UT7 cells (Figure 4D). Therefore, these results strongly suggest that miR-495-3p also modulates Ponatinib sensitivity in Imatinib-resistant UT7-T315I cells.

**Figure 4:**
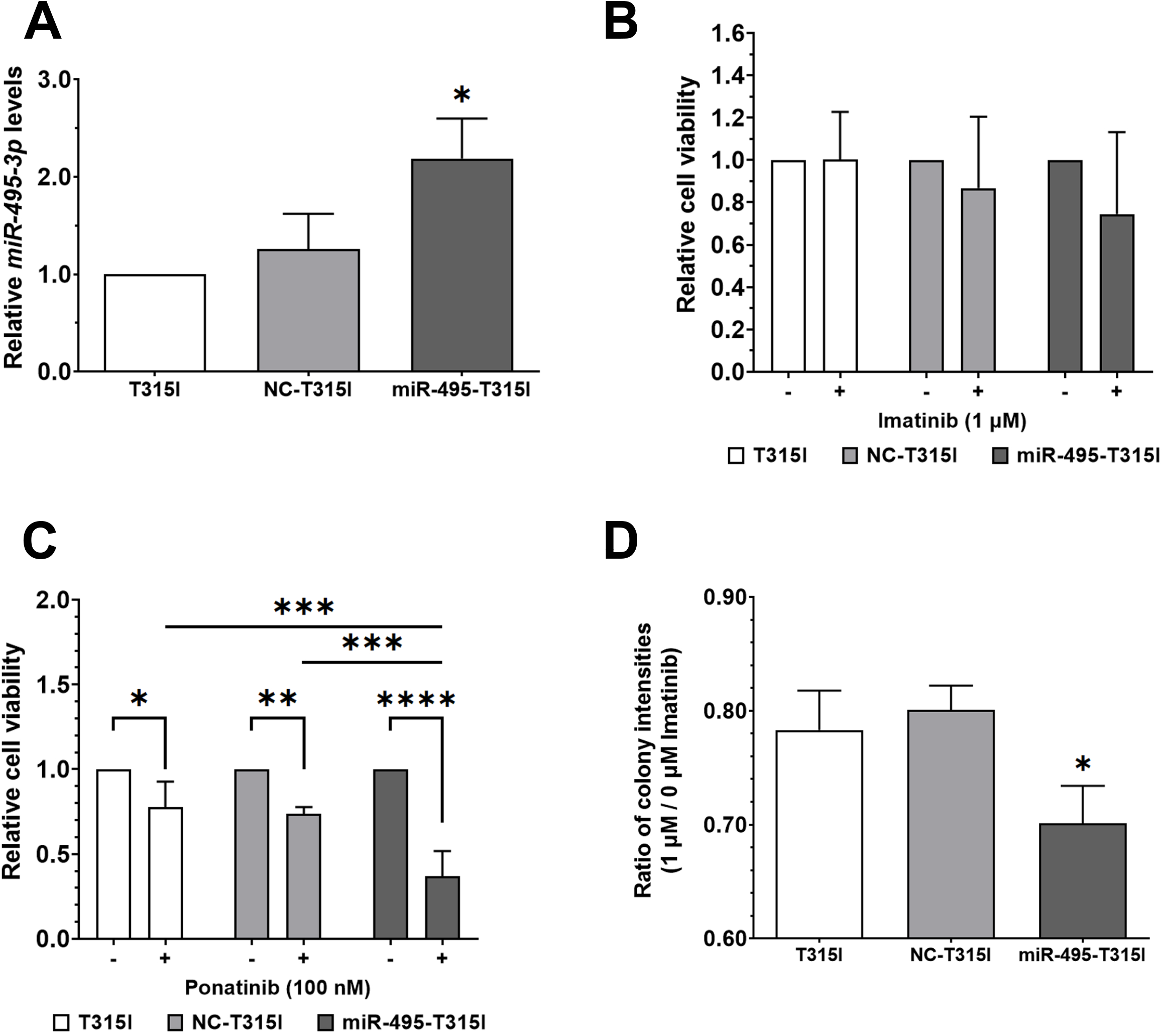
Overexpression of *miR-495-3p* in TKI-resistant CML cells. **(A)** Relative quantification of *miR-495-3p* expression by real-time PCR of UT-7/11 T315I cells stably expressing miR-495-3p (miR-495-T315I) or its control vector (NC-T315I) (FC = 2.19 ± 0.41, p = 0.0210). *miR-495-3p* expression was normalized to *U6* expression. **(B-C)** Cytotoxic effects of IM (B) or Ponatinib (C) were measured using XTT assay (22.39% reduction in UT7-T315I, 26.31% in NC-T315I UT7, and 63.00% in miR-495-T315I, p=0.0003 compared to UT7-T315I and p=0.0008 compared to NC-T315I). **(D)** Cytotoxic effect of Ponatinib on colony formation of miR-NC or miR-495-3p overexpressing cells. CFC assays were performed in the absence [0 μM] or presence of 100 nM Ponatinib (mean ratio = 0.70 ± 0.03 versus 0.78 ± 0.03 in UT7-T315I cells, p=0.0372). The bar chart shows the difference in the ratio of colony intensity as a quantitative parameter of IM treatment efficiency. Each data represents mean ± SD from three independent experiments. The One-way or Two-way ANOVA approach was used to assess the significance. *, p < 0.05; **, p < 0.01; ***, p < 0.001; ****, p < 0.0001.

### Bioinformatics analyses of the miR-495-3p targets during CML progression and identification of molecular partners

To investigate whether *miR-495-3p* could play a role during CML progression, we analyzed the expression of its target genes in CD34+ cells from CP and BC patients using bioinformatics tools. Putative targets were predicted through miRDB and Targetscan human databases with a common overlap comprising 466 transcripts, including 366 that could be analyzed in GSE4170 dataset comprising transcriptome from patients at different CML stages (Figure 5A). Interestingly, unsupervised principal component analysis based on the expression of the 366 spotted mRNA targets allowed a clear separation between CML samples from blast crisis and chronic phase (Figure 5B). Strikingly, differential expressed gene analysis between CP *versus* BC identified a set of *miR-495-3p* target genes up-regulated during the advanced phase of the disease (Figure 5C). Indeed, the up-regulation of these target genes allowed to stratify hematopoietic cells from BC and CP (Figure 5D). Among these target genes, several have already been described as upregulated during BC such as *TM4SF1, ZNF827, KLHL5*, and *CTNND2* [32,41], or implicated in leukemia pathophysiology *such as GAB1, LHX2, THRB*, and *CDH2* [42–45] (Figure 5D). Next, we built a protein-protein interaction network using this gene set. Based on blood differentiation gene interaction, this protein-protein interaction network collected 973 edges between 872 nodes from 32 target gene seeds (Figure 5E). Functional inference of the Gene Ontology Biological Process database on the network highlighted a significant implication of 211 molecular partners implicated in protein phosphorylation (Figure 5E). Altogether, these results suggest that during BC, a set of *miR-495-3p* target genes are up-regulated in hematopoietic progenitors and recruit partners that are implicated in the protein phosphorylation cell functionality.

**Figure 5:**
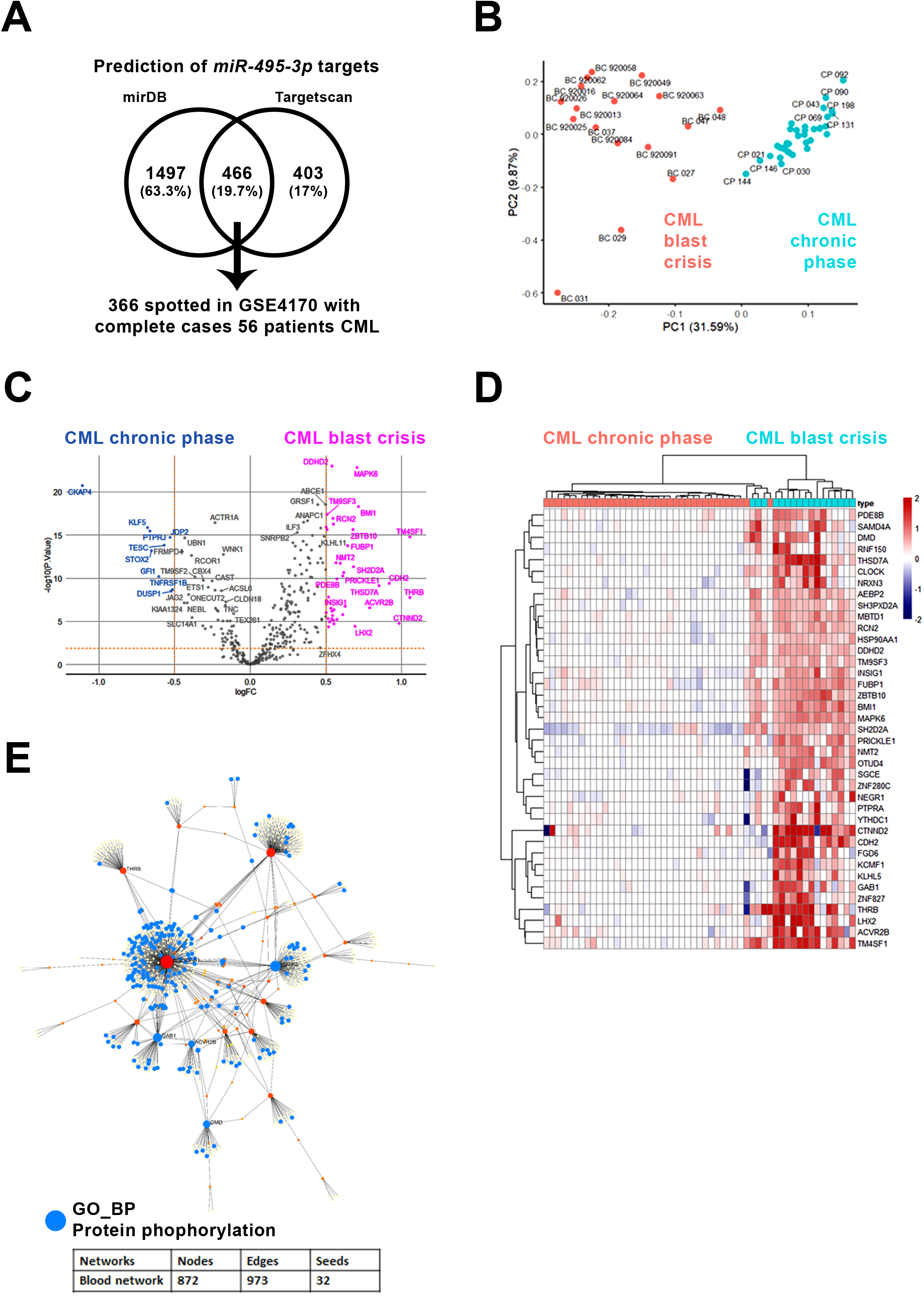
miR-495-3p regulates targets affecting protein phosphorylation during CML blast crisis. **(A)** Venn diagram of miR-495-3p target prediction overlap and quantified in GSE4170 transcriptome dataset. **(B)** Unsupervised principal component analysis of miR-495-3p targets in CD34+ hematopoietic progenitors from CML patients in chronic *versus* blast crisis phase. **(C)** Volcano-plot representing the differential expression of predicted miR-495-3p target genes between CML patients in blast crisis *versus* chronic phase chronic. **(D)** Heatmap representing miR-495-3p targets significantly upregulated in CD34+ hematopoietic progenitors obtained from CML patients in blast crisis *versus* chronic phase **(E)** Schematic representation of protein-protein interaction network based on miR-495-3p targets found up-regulated during CML blast crisis (p = 3.69e-42). Functional inference on the network revealed in blue a main implication of protein phosphorylation.

### Evaluation of the expression of *miR-495-3p* target genes in AML

We next analyzed the expression status of *miR-495-3p* target genes up-regulated in BC using RNA-sequencing transcriptome from AML TCGA cohort. Among these target genes, 8 were found strictly overexpressed in the blasts of AML patients, including *BMI1*, SH2D2A, *INSIG1, HSP90AA1, LHX2, NRXN3, SH3PXD2A*, and *GAB1* (Figure 6A). The combined alterations affecting these 8 target genes in blast AML allowed to stratify two groups of AML patients herein referred as altered (associated with the overexpression of *miR-495-3p* target genes) and unaltered survivals. Overall survival analysis performed on this patient stratification identified a significantly worse prognosis in the altered group as compared to the unaltered group (Figure 6B). Interestingly, this stratification also identified that the altered group of patients exhibits a higher level of genomic instability and a reduced number of platelets (Figure 6C, 6D). Taken together, these results suggest that overexpression of *miR-495-3p* target genes is also associated with a dismal prognosis in AML as compared to the group with low expression of *miR-495-3p*.

**Figure 6:**
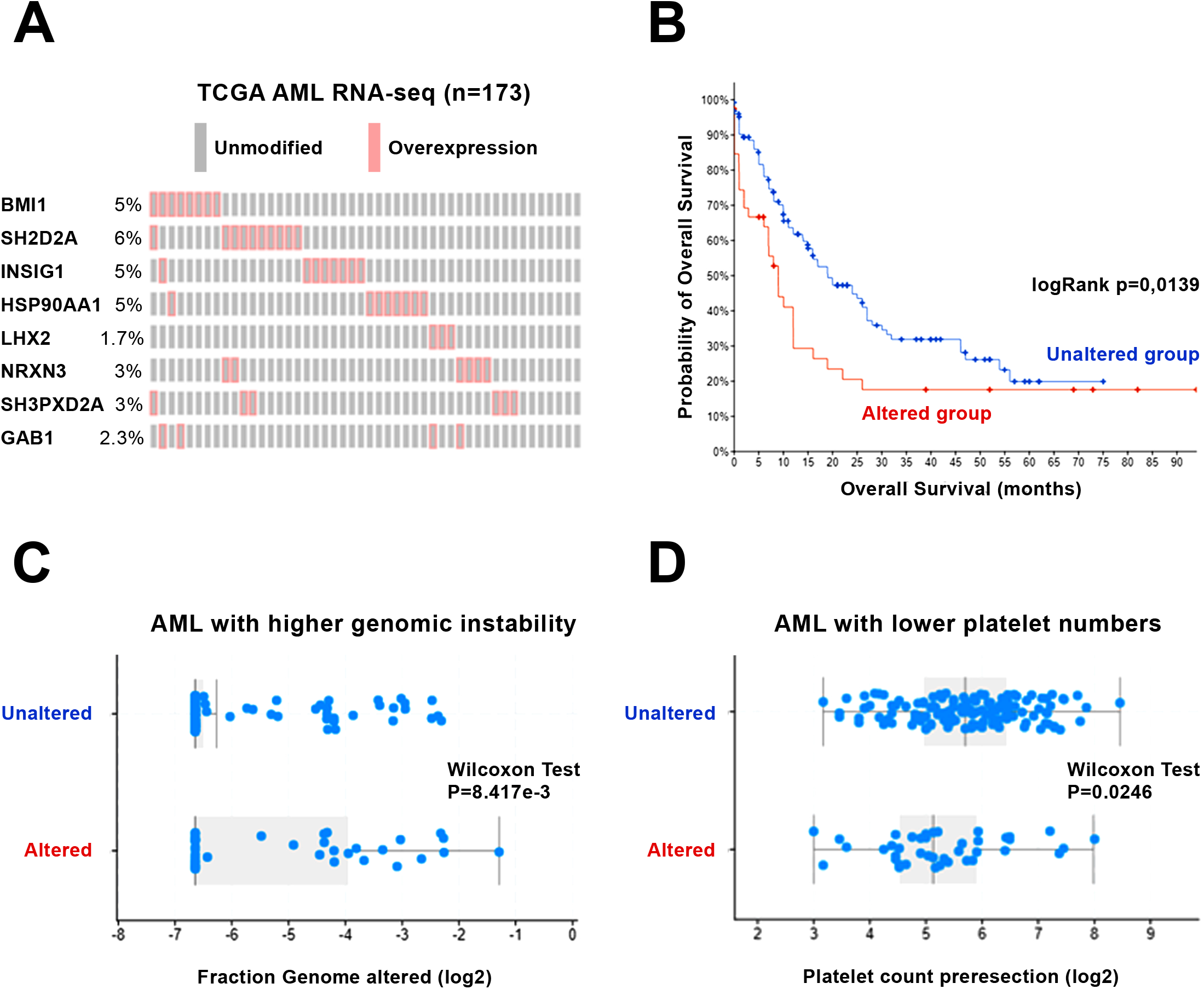
Up-regulation of miR-495-3p targets defined a worse prognosis in AML. Analyses were performed on omics experiments of the TCGA AML cohort. **(A)** Oncoprint visual output of the expression of miR-495-3p target genes in AML leukemic cells (red square, high expression; grey square, no alteration). Percentages indicate the proportion of the patients. **(B)** Overall survival Kaplan-Meier curves stratified on combined transcriptional alteration of miR-495-3p target genes in AML leukemic cells (log rank p = 0.0139). **(C)** Quantification of the genomic instability stratified on combined transcriptional alterations of miR-495-3p targets in AML leukemic cells (p = 8.417e-3). **(D)** Quantification of the platelet number in AML patients stratified on combined transcriptional alterations of miR-495-3p targets in AML leukemic cells (p = 0.0246).

## DISCUSSION

Over the last two decades, microRNAs have emerged as important regulators of cancer progression and drug resistance. In CML, microRNAs can exert a tumor suppressor activity by directly targeting *BCR::ABL1*, cell cycle regulators, and hematopoiesis regulators. Alternatively, they can also exhibit oncogenic properties by blocking for example myeloid differentiation or targeting tumor suppressor genes [21].

In this study, we uncovered for the first time a role of *miR-495-3p* in BCR::ABL1 expressing leukemic cells. We first observed a reduced expression in BCR::ABL1 expressing cell lines. Interestingly, using a *BCR::ABL1* inducible UT-7 cell line, we observed a downregulation of *miR-495-3p* expression when BCR::ABL1 expression was induced, suggesting that BCR::ABL1-mediated signaling pathways might regulate microRNA expression (Figure 1B). This could potentially be associated with the results of our bioinformatics analyses showing upregulation of *miR495-3p* putative target genes in patients in BC compared to the ones in CP, suggesting that miR-495-3p activity is reduced during CML progression. These findings however remain to be validated in independent cohorts of patients. Similarly, the regulation of the expression of *miR-495-3p* from a mechanistic view will require further investigation. Particularly, *miR-495-3p* expression has been linked to DNA methylation [46], the expression of E12/E47 transcription factors [47], or oncogenic fusion protein such as MLL-AF9 [48]. As an alternative but not exclusive hypothesis, the downregulation of *miR-495-3p* expression in the bone marrow of the patients could reflect a change in the cellular composition of the bone marrow with the dramatic increase of immature blast cells at advanced stages of the disease. Particularly, it is tempting to speculate that *miR-495-3p* expression reflects the number of immature blastic cells which could be used as a prognosis marker.

In previous studies, *miR-495-3p* was found to be downregulated in AML and a higher expression was associated with a worse prognosis [49,50]. Similarly, *miR-495-3p* expression is downregulated and acts as a tumor suppressor in MLL-rearranged leukemia [48]. These observations suggest that *miR-495-3p could* act as a tumor suppressor. To gain more insights into the function of this microRNA, we undertook the analysis of the phenotypes resulting from both complementary gain- and loss-of-function strategies in BCR::ABL1-expressing cell lines.

Our first observation is that miR-495-3p expression is low in cells expressing BCR::ABL1. This phenomenon seems to be dependent on the expression of BCR::ABL1 as shown in the DOX-inducible BCR::ABL1 cell in which DOX induces the transcriptional activation of BCR::ABL1. On the other hand, AntagomiR-495 stimulates cell proliferation while miR-495-3p mimics induced a reverse phenotype associated with an increase of cells in G0/G1 phase. We propose a model in which *miR-495-3p* downregulation during CML progression might contribute to the expansion of immature blast cells. It has to be stressed that the cell lines in which these results have been obtained are essentially close to BC samples. Interestingly, *miR-495-3p* was previously described as a negative regulator of cell proliferation in several solid tumors by targeting *CDK6* in glioblastoma multiforme [51]. Inversely, *miR-495-3p* promotes cell proliferation in breast cancer presumably through the release of mTOR activity [47]. Further experiments will be required to clarify how *miR-495-3p* regulates cell proliferation in CML cells.

Our second observation is that miR-495-3p promotes TKI resistance. Indeed, UT-7/11 cells expressing AntagomiR-495 exhibit an increased clonogenic survival in presence of IM (Figure 1H) while the phenotype is reversed when *miR-495-3p* is overexpressed (Figure 2F). We proposed that these phenotypes are linked to *MDR1* expression which encodes for a major actor of multidrug resistance. Interestingly, *miR-495-3p* was previously identified as a direct negative regulator of *MDR1* expression in a cellular model of multidrug resistance induced in CML K562 cells by Adriamycin or Vinblastin [25]. We thus extend this important role of miR-495-3p in drug resistance to IM which is a major TKI used in clinics. Strikingly, in a model of TKI resistance induced by T315I mutation of BCR::ABL1, *miR-495-3p* overexpression did not alter their sensitivity to IM but to Ponatinib. Whether similar mechanisms involving MDR1 are at play remains to be determined in the context of Ponatinib. Indeed, Ponatinib was previously shown to be transported independently from MDR1 in K562 [52]. MDR1 overexpression in K562 cells confers resistance to IM but to a weaker extent to Ponatinib [53]. Alternatively, miR-495-3p could regulate drug resistance through other mechanisms. For example, miR-495-3p targets *ATP7A* and *UCA1/NRF2* in non-small cell lung cancer [54,55], and *TSPAN12* in small cell lung cancer [56].

Taken together, our results strongly suggest that miR-495-3p plays a role in the proliferation and drug resistance of BCR::ABL1-expressing leukemic cells. The expression of miR-495-3p targets is associated with CML progression. These findings open new avenues in the development of treatment and the identification of biomarkers that could be used to better stage CML and predict responses to TKI treatment. Indeed, *miR-495-3p* could be used as a biological marker for prognosis and/or pharmaceutical target.

## ACKNOWLEDGEMENTS

We are very grateful to the flow cytometry core facility of UMS-44. We appreciate expert technical supports by Dr. Hervé Acloque. This work was supported by the ATIP/AVENIR INSERM program and Paris Saclay. Y.R. received a Franco-Thai scholarship from the Embassy of France in Bangkok and funding from Vaincre le Cancer NRB. I.S. received funding from the Ministère de l’Enseignement Supérieur de la Recherche et de l’Innovation.

